# Effect of chronic upregulation of endocannabinoid signaling *in vivo* with JZL184 on striatal synaptic plasticity and motor learning in YAC128 Huntington disease mice

**DOI:** 10.1101/2024.09.25.614804

**Authors:** Marja D. Sepers, Cameron L. Woodard, Daniel Ramandi, Haley A. Vecchiarelli, Matthew N. Hill, Lynn A. Raymond

## Abstract

Synaptic dysfunction underlies early sensorimotor and cognitive deficits, and precedes neurodegeneration in a variety of disorders, including Alzheimer, Parkinson and Huntington disease (HD). A monogenic inherited disorder, HD manifests with cognitive, motor and mood disorders associated with progressive degeneration of striatal spiny projection neurons and cortical pyramidal neurons. Cortico-basal ganglia-thalamic loops regulate movement selection and motor learning, which are impaired early in HD. Skilled motor learning is mediated in part by plasticity at cortico-striatal synapses, including endocannabinoid-mediated, high-frequency stimulation induced long-term depression (HFS-LTD). Previously, we found impaired HFS-LTD in brain slice recordings from pre-manifest HD mouse models, which was corrected by JZL184, an inhibitor of endocannabinoid 2-arachidonoyl glycerol (2-AG) degradation. Here, we tested the effects of JZL184 administered *in vivo* to YAC128 HD model and wild-type (WT) littermate mice. JZL184, given orally daily over a 3-week period, significantly increased levels of 2-AG in striatal tissue. While JZL184 treatment had no impact on open field behavior which was similar for the two genotypes, the treatment improved motor learning on the rotarod task in YAC128 mice to the level observed in WT mice. Moreover, HFS-induced striatal plasticity measured by field potential recording in acute brain slice from YAC128 mice was normalized to WT levels after JZL184 treatment. These results suggest a novel target for mitigating early symptoms of HD, and support the need for clinical trials to test the efficacy of modulating the endocannabinoid system in treatment of HD.

## INTRODUCTION

Cortical-basal ganglia-thalamic loops are involved in the control of movement (Bariselli et al., 2019; Krack et al., 2010), and plasticity at synapses between glutamatergic cortical afferents and GABAergic spiny projection neurons (SPNs) of the dorsal striatum is thought to be involved in motor learning (Dayan and Cohen, 2011; Kreitzer and Malenka, 2008; Perrin and Venance, 2019). Studies that have measured neuronal activity in motor cortex and dorsal striatum while mice learn to run on an accelerating rotarod have shown that neurons in both structures have dynamic changes in their firing activity over the course of days of repeated trials (Costa et al., 2004; Kupferschmidt et al., 2017; Yin et al., 2009). Modulatory, G-protein coupled receptors and their downstream signaling pathways have been implicated when synaptic plasticity is induced by trains of cortical stimuli in *ex vivo* brain slice recordings. One prominent mechanism is endocannabinoid (eCB)-dependent long-term depression (LTD) in striatal SPNs (Lovinger, 2010), which occurs primarily at corticostriatal but not thalamostriatal synapses (Wu et al., 2015). eCBs 2-arachidonoylglycerol (2-AG) and N-arachidonoylethanolamine (AEA or anandamide) are lipid mediators that are synthesized in the postsynaptic compartment and diffuse to the presynaptic membrane where they bind cannabinoid 1 receptor (CB1R) to down-regulate release of fast neurotransmitters glutamate and GABA (Araque et al., 2017; Lovinger, 2010). CB1R is highly expressed in the dorsal striatum, including on cortical terminals; high frequency stimulation (HFS) of these afferents induces eCB synthesis in striatal SPNs, resulting in suppression of presynaptic glutamate release and long-term depression (LTD) at these corticostriatal synapses (Lovinger, 2010). eCB-dependent LTD is thought to be involved in motor learning (Araque et al., 2017; Lovinger, 2010).

Huntington disease (HD), which is the most common hereditary neurodegenerative disorder, is caused by a CAG triplet repeat expansion in the *HTT* gene (“A novel gene containing a trinucleotide repeat that is expanded and unstable on Huntington’s disease chromosomes. The Huntington’s Disease Collaborative Research Group,” 1993). HD manifests as a disorder of movement, cognition and mood, and SPNs in the dorsal striatum are the most severely affected by neurodegeneration, along with cortical pyramidal neurons (CPNs) (Bates et al., 2015; Ross and Tabrizi, 2011). In carriers of the *HTT* mutation, skilled motor learning is often impaired before a clinical diagnosis (Doyon, 2008; Feigin et al., 2006; Holtbernd et al., 2016). Mouse models of HD facilitate understanding mechanisms underlying early synaptic changes that precede neurodegeneration (Cepeda and Levine, 2022; Milnerwood and Raymond, 2010; Raymond et al., 2011), including deficits in cortical-striatal synaptic plasticity (Plotkin et al., 2014; Sepers et al., 2018). Prior to motor manifestations in 2 month-old YAC128 HD mice, we previously recorded from *ex vivo* cortical-striatal slices and reported altered striatal SPN glutamate receptor function (Milnerwood et al., 2010; Milnerwood and Raymond, 2007) and a deficit in cortico-striatal HFS-induced LTD that worsens with disease stage (Sepers et al., 2018). The latter is a result of impaired synthesis of the eCB anandamide, but HFS-LTD can be restored by treatment with JZL184, an inhibitor of monoacylglycerol lipase (MAGL), which degrades 2-AG (Sepers et al., 2018).

Here, we tested the effect of orally administered JZL184 on motor learning on the accelerating rotarod task, performance in the forced swim test and open field, and *ex vivo* corticostriatal synaptic plasticity, in YAC128 mice compared to WT littermates. Results suggest that modulation of endocannabinoids is a promising strategy for treatment of early motor manifestations in HD.

## METHODS

### Animals and Experimental Design

All procedures were carried out in accordance with the Canadian Council on Animal Care and approved by the University of British Columbia Committee on Animal Care (protocols A19-0076, A23-0083). Experiments used male heterozygous YAC128 transgenic mice (*n* = 20; line 53; Slow et al., 2003) on the FVB/NJ background, with male wild-type (WT) littermates as controls (*n*=18). Male mice were used for these experiments to match the cohort of male mice that showed attenuated eCB-mediated HFS-LTD in cortical-striatal brain slice (Sepers et al., 2018). Mice of both genotypes were housed together and were 5-months-old (+/- 2 weeks) at the start of treatment. Animals were housed on a 12/12 h light/dark cycle in a temperature and humidity-controlled room and given environmental enrichment (i.e. hut, PVC tubing) and *ad libitum* access to food and water. Tissue from ear-clipping was used for DNA extraction and PCR analysis to determine genotype.

The experimental design is shown in Fig. 1A. Prior to the start of experiments, WT and YAC128 mice were randomly assigned to treatment and vehicle control groups, and handled on at least two occasions for 3 minutes per animal. Mice were treated at the same time each day with JZL184 (4mg/kg, oral) or vehicle, and a battery of behavioral tests was performed from Day 15 (D15) to D20 of treatment. Behavioral testing began approximately two hours following treatment on each day. Tissue collection for endocannabinoid analysis and slice electrophysiology was performed from D21 to D25 of treatment, two to six hours following final drug administration. The behavioral testing and electrophysiology was done at a time when JZL184 was active and elevating brain tissue levels of 2-AG (see Fig. 1B) in order to mimic conditions of a clinical trial in patients with HD. Drug treatment and behavioral testing was performed during the light phase of the light cycle. The experimenter was blinded to both genotype and treatment condition throughout testing.

**Figure 1.**
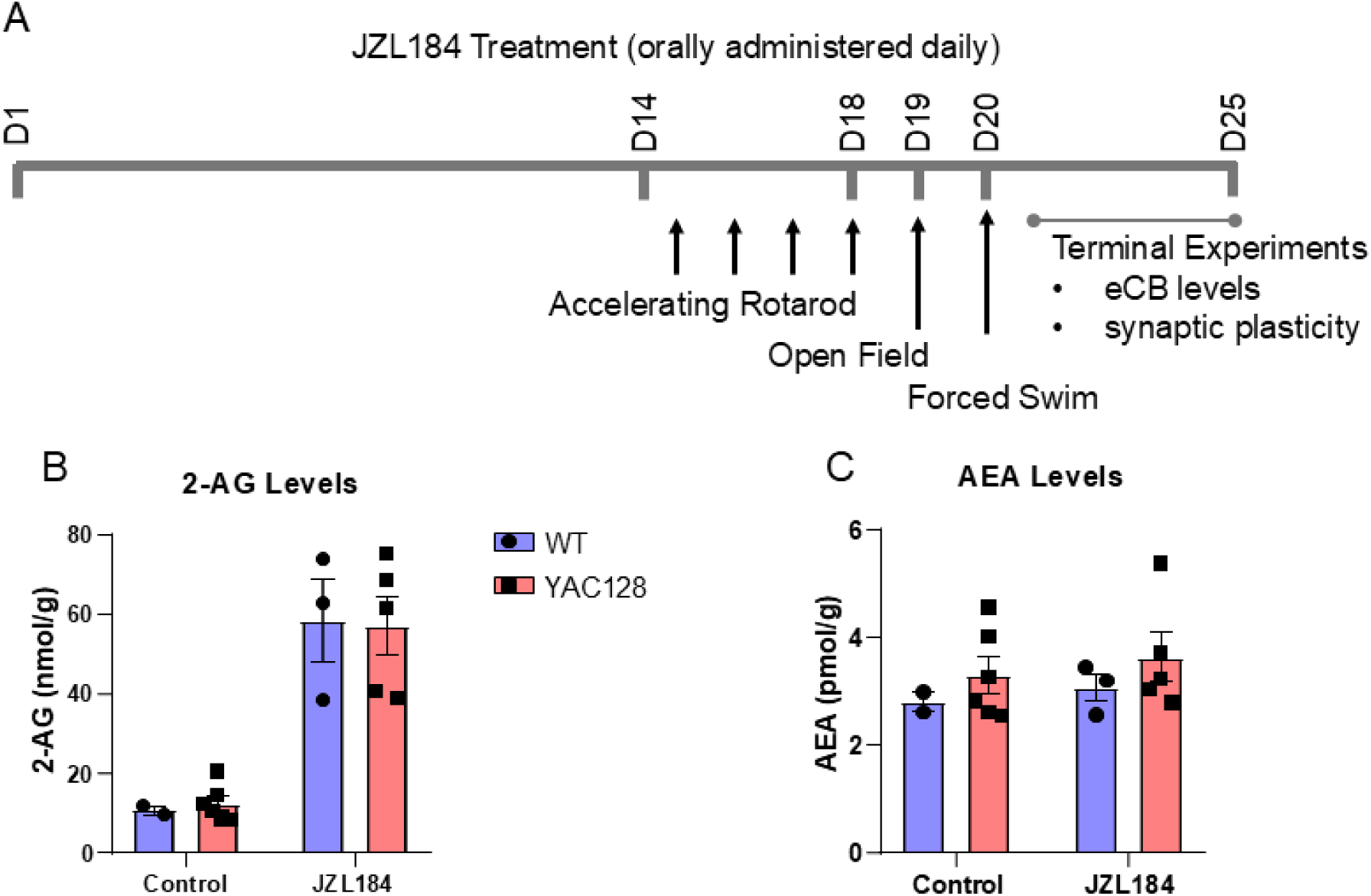
Chronic oral treatment with JZL184 elevates striatal 2-AG. A) Timeline of JZL184 drug treatment experiments in 5-month-old YAC128 HD mice and WT littermates. B) 2-AG levels in WT and YAC128 mice were significantly increased by JZL184 treatment (*p* < 0.0001). C) Anandamide (AEA) levels were unchanged following treatment. Each point represents measurement from striatal tissue of one mouse.

### Drug Treatment

To allow for voluntary oral drug dosing, JZL184 (Tocris 3836) was dissolved into peanut butter as done previously (Doenni et al., 2016; Long et al., 2009; Qi et al., 2015). Briefly, peanut butter was warmed in a double boiler on a hot plate while stirring. Drug was added at a ratio of 2 mg of JZL184 per 1 gram of peanut butter and mixed for 15 minutes. Peanut-butter-drug mixture was aliquoted and frozen until just prior to administration. Mice were given plain peanut butter once per day for three days prior to start of treatment to habituate them to eating peanut butter. On treatment days, mice were weighed and given 2 mg of peanut-butter-drug mixture per gram of bodyweight, for an effective dose of 4mg/kg JZL184 per day. Vehicle-treated mice were given an equivalent amount (2 mg/g bodyweight) of plain peanut butter. Peanut butter was administered in a plastic dish at the same time each day in a separate cage from the home cage, and mice were monitored to ensure that they consumed the full amount. Drug dosage, frequency, and length of treatment was determined based on previous literature (Feliszek et al., 2016; Ghosh et al., 2015; Long et al., 2009; Schlosburg et al., 2010) and preliminary dosing studies in our lab.

Additional cohorts of WT mice were used for electrophysiological experiments. Naive mice were not handled or given peanut butter (*n* = 3). Mice without behavioral testing received peanut butter as above with daily handling (PB only) before making striatal slices but were not tested on rotarod, open field or forced swim (*n* = 5).

### Behavioral Testing

#### Accelerating rotarod test

To measure motor coordination and learning, mice were assessed for 4 consecutive days on the accelerating rotarod test (D15-D18 of treatment). Mice are placed on a rotating rod that accelerates from 5 - 40 RPM in 300 s, and they must continuously adjust their position in order to avoid falling off. Latency to fall was recorded when the mouse fell off, or else completed a full rotation while holding onto the rod, before the maximum time of 300 s. If the mouse reached the maximum time without falling, the trial was ended and scored as 300 s. Each mouse performed 3 trials per day, for a total of 12 trials. Testing was performed at the same time each day.

#### Open field test

The open field test was used to assess locomotor activity levels. Mice were placed one at a time in the corner of an open-top clear acrylic box (48 × 38 cm) under bright lighting and were allowed to explore for 10 minutes. Open field activity was recorded at 30 frames per second by a camera (Waveshare, 10299) mounted above the box. DeepLabCut (Mathis et al., 2018) was used to track the position of the mouse in each video, and total distance traveled was quantified using custom MATLAB scripts. Time spent in the central zone of the open field (33 × 23 cm) was also quantified. Open field video recordings were lost for one cohort of mice (*n =* 13) due to hard drive failure and could not be analyzed.

#### Forced swim test

The forced swim test was used to assess coping responses to stressful stimuli. Mice were placed one at a time in a large beaker (30 cm height x 20 cm diameter) filled with 15 cm of warm (24-25°C) water, so that the animal cannot touch the bottom of the beaker or escape from the top and has to swim or float for the duration of the trial. Activity was monitored for a period of five minutes and all movements were recorded on video. The water in the beaker was changed between each animal. An observer (blinded to genotype and treatment) scored the videos by measuring the amount of time the mouse spent floating during the last 4 minutes of the trial. Forced swim testing was performed last to prevent any possible confounds of swim stress on subsequent behavior.

### Mass Spectrometry

Concentrations of eCB were measured as previously described (Qi et al., 2015). Briefly, glass tubes containing 2 mL of acetonitrile and 100 µL of IS (AEA-d8 at 0.1 μM and 2-AG-d8 at 5 μM) were prepared for brain samples, which were weighed and the frozen piece placed into the prepared glass tubes for manual homogenization with a glass rod until resembling sand. Samples were then sonicated, stored overnight at −20°C to precipitate proteins, centrifuged (1800 rpm at 4°C for 3-4 min) to remove particulates, and the supernatant transferred to a new glass tube. Tubes were then placed under nitrogen gas to evaporate, sidewalls washed with 250 µL acetonitrile to recollect any adhering lipids, evaporated again, and re-suspended in 100 µL of acetonitrile into a glass vial with a glass insert. Resuspended samples were stored at −80°C until analysis by LC-MS/MS. Analyte (AEA and 2-AG) concentrations (in pmol and nmol/µL respectively) were normalized to sample weight. The lower limit of quantification for AEA was 0.035 ng/ml and for 2-AG was 19 ng/ml; the upper limit of quantification for AEA was 35 ng/ml and for 2-AG was 18950 ng/ml.

### Electrophysiology

For electrophysiology experiments, mice were anesthetized with isoflurane and decapitated. Sagittal brain slices 250μm thick containing the striatum were made on a vibratome (Leica VT100) in ice-cold artificial cerebrospinal fluid (aCSF) with (in mM): 125 NaCl, 2.5 KCl, 25 NaHCO_3_, 1.25 NaH_2_PO_4_, 10 glucose, 0.5 CaCl_2_ and 2.5 MgCl_2_. Slices were then transferred into a holding chamber at 37°C for 30 - 45 min. In the holding chamber and for all experiments the aCSF was made as above but contained 2mM CaCl_2_and 1mM MgCl_2_ with pH of 7.3, osmolarity 310 mosmol and oxygenated with 95%O_2_ / 5% CO_2_. Slices were maintained at room temperature for at least 2 hours before being transferred to the recording chamber and continuously superfused at room temperature with aCSF containing 50μM picrotoxin (Tocris). Extracellular field recordings were made with a glass micropipette containing aCSF placed in the dorsal striatum with a glass stimulating pipette positioned more than 200uM away. Test stimuli were delivered every 15 seconds and field excitatory postsynaptic potentials (fEPSP) recorded using an Axopatch 200B or Multiclamp 700 amplifier and Clampex 10.7 software (Molecular Devices), sampled at 100kHz and filtered at 1kHz.

### Statistics

All statistical testing was performed using Prism 8 (GraphPad Software). Behavioral data was analyzed using regular or repeated measures two-way ANOVA followed by Sidak’s multiple comparisons test. Alpha level for all tests was *p* = 0.05. For behavioral and mass spectrometry experiments, *n=* the number of animals.Three mice were identified post-hoc who had very high body weight (50-60 grams at start of testing). These mice were removed from analysis for rotarod testing, as high bodyweight is a known confound for rotarod testing (McFadyen et al., 2003) and these mice often fell off immediately after being placed on the rod. One mouse was removed from analysis for the forced swim test due to being identified as an outlier using the ROUT method (Q = 1%). For electrophysiology experiments, *n=* the number of slices and animal numbers are given in brackets. Cumulative response to HFS was analyzed by repeated measures ANOVA with Sidak’s multiple comparisons. To categorize plasticity, raw fEPSP baseline amplitudes were compared to amplitudes 30min after HFS for each experiment by t-test.

## RESULTS

Previously, we reported that LTD of excitatory synaptic responses recorded from striatal SPNs in response to high-frequency cortical stimulation in brain slice (HFS-LTD) is impaired early in YAC128 mice compared with WT littermates (Sepers et al., 2018). We and others have shown this form of cortico-striatal synaptic plasticity is a result of reduced probability of glutamate release from presynaptic cortical terminals, mediated by CB1R activation from the release and retrograde diffusion of endocannabinoids from striatal SPNs (Koch et al., 2018; Lovinger, 2010; Sepers et al., 2018). Our study also showed that although anandamide synthesis, which is normally stimulated by HFS and involved in mediating this form of LTD (Lerner and Kreitzer, 2012), was impaired, HFS-LTD could be restored in YAC128 brain slice by treatment with JZL184 to inhibit degradation of 2-AG (Sepers et al., 2018). Since corticostriatal synaptic HFS-LTD has been implicated in motor learning (Araque et al., 2017; Lovinger, 2010; Yin et al., 2009), in this study we set out to determine whether treatment with JZL184 could restore HFS-induced synaptic plasticity, improve motor learning or impact performance on other behavioral tasks in YAC128 mice.

### Oral treatment with JZL184 elevates striatal 2-AG but not anandamide levels

We treated 5-month old YAC128 mice and their littermate wild-type controls for up to 25 days with 4mg/kg/day JZL184 mixed in peanut butter, or peanut butter alone (vehicle-treated). The treatment and behavioral testing timeline is shown in Fig. 1A. Between days 20 - 25 after starting treatment, mice were sacrificed for terminal *ex vivo* brain slice field recording from the striatum of one hemisphere, and striatal tissue from the opposite hemisphere was extracted and flash frozen on dry ice. Samples of frozen tissue were subjected to lipid extraction and analysis via mass spectrometry, as described above (see Methods). As expected, after 20-25 days of treatment with JZL184, striatal 2-AG levels were increased more than 5-fold above baseline (RM ANOVA, treatment: *F*(1, 23) = 41.13, *p <* 0.0001) (Fig. 1B), whereas anandamide levels were unchanged (RM ANOVA, treatment: *F(*1, 23) = 0.53, *p =* 0.48) (Fig. 1C). No differences were observed in baseline levels of eCBs or in the effect of treatment on WT versus YAC128 mice.

### Rotarod performance improved in YAC128 mice with JZL184 treatment

Previous studies have suggested that HFS-induced synaptic plasticity at cortico-striatal synapses, as recorded from acute slice, contributes to motor learning *in vivo* (Balleine and O’Doherty, 2010; Graybiel and Grafton, 2015; Klaus et al., 2019; Perrin and Venance, 2019). To determine the effect of increasing 2-AG levels with JZL184 on motor learning, we tested mice on an accelerating rotarod (5 - 40 rpm over 5 minutes). Consistent with previous studies (Menalled et al., 2012), we found that vehicle-treated YAC128 mice performed significantly worse than WT mice in learning to stay on the rotarod (RM ANOVA, genotype: *F*(1, 15) = 6.62, *p* = 0.02) (Fig. 2A), although both genotypes improved their performance across testing (day: *F*(3, 45) = 21.4, *p* < 0.0001). Latency to fall from the rotarod on the last day of testing was significantly lower in YAC128 mice as compared to WT (Sidak, *p* = 0.04) (Fig. 2C) In contrast, after treatment with JZL184, YAC128 mice performed equally well to treated WT mice on the rotarod test (RM ANOVA, genotype: F(1, 16) = 0.6, p = 0.45) (Fig. 2B,C).

**Figure 2.**
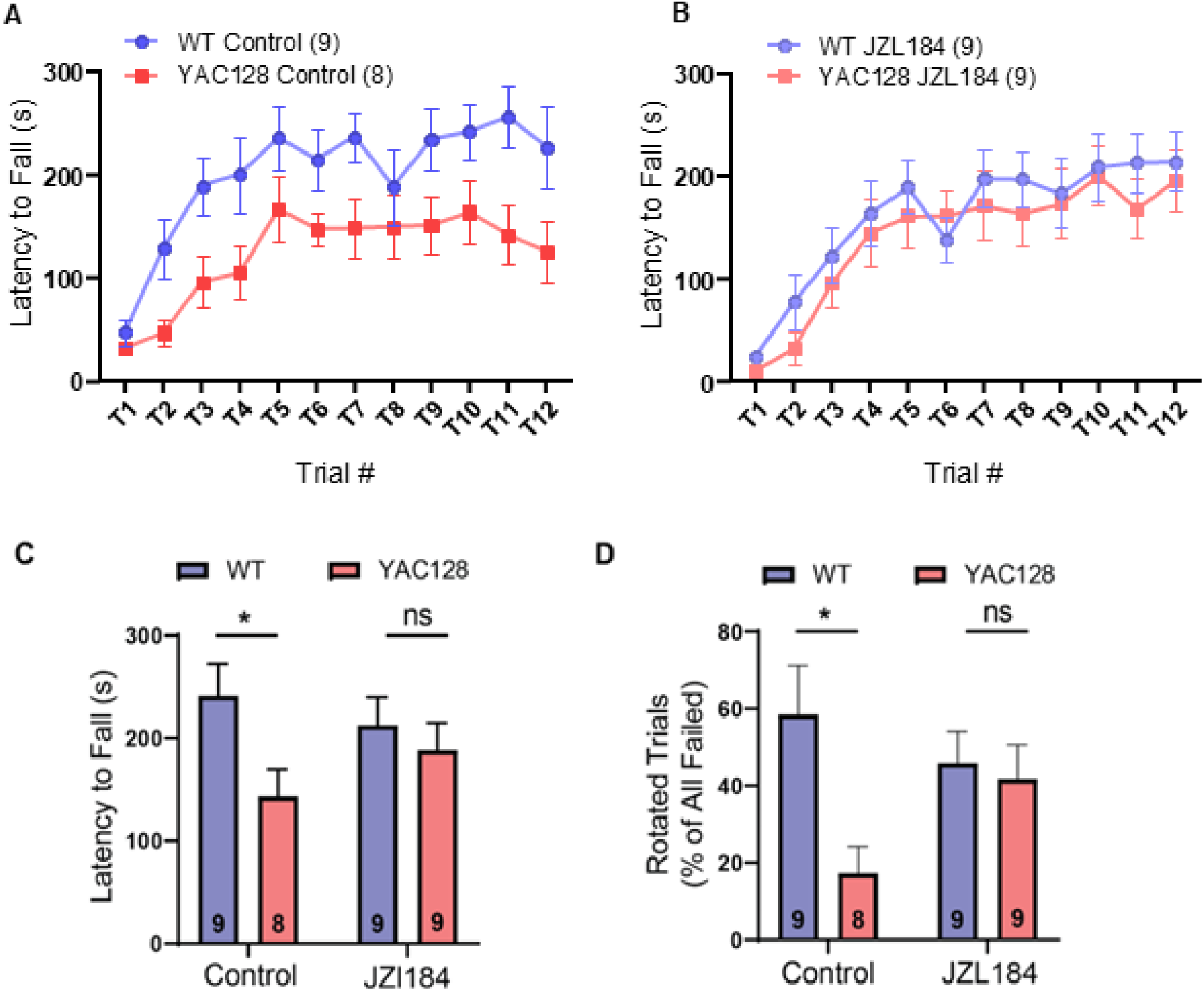
Motor coordination assessed using the accelerating rotarod test. A) Control (vehicle treated) YAC128 mice showed impaired performance on the rotarod compared to WT mice (*p =* 0.02). B) Following JZL184 treatment, rotarod learning was not significantly different between WT and YAC128 mice. C) Latency to fall off the rotarod on day 4 (last day of testing) was significantly lower in control, but not JZL184-treated YAC128 mice. D) Failed trials ended when the mouse fell or rotated around the rod. Fewer trials ended by rotation in control, but not JZL184-treated mice as compared to WT. \**p*<0.05 by Sidak’s test. Numbers in parentheses (A,B) and in bars (C,D) represent number of mice.

Both a full rotation while holding onto the rod and a direct fall from the rod were considered trial endpoints for the rotarod task. Control Vehicle-treated YAC128 mice had significantly fewer trials that ended in rotations as compared to WT mice (Sidak, *p* = 0.01) (Fig. 2D). Interestingly, for JZL184-treated YAC128 mice, in addition to the marked improvement in overall latency to fall over the four days, the percentage of trials that ended with rotations versus falls also increased ~2-fold and was not significantly different from WT (Sidak, *p =* 0.94), consistent with improved coordination and/or strength (Fig. 2D).

### JZL184 treatment increases immobility in forced swim test but does not change open field performance

Studies have reported that JZL184-mediated elevation of 2-AG can affect coping responses to stress and anxiety-like behaviors in rodents (Busquets-Garcia et al., 2011; Pavón et al., 2021). HD mouse models have been shown to exhibit increased immobility in the forced swim test (FST) (Pouladi et al., 2013; Southwell et al., 2016), which is thought to test stress reactivity and passive versus active coping responses (Pavón et al., 2021). Here, we compared WT and YAC128 mice treated with JZL184 or vehicle in the FST to determine if elevating 2-AG levels could affect this phenotype in HD mice. As expected, YAC128 mice showed increased immobility time in the FST compared with WT mice (ANOVA, genotype: *F*(1, 33) = 4.28, *p* = 0.046) (Fig. 3A). Strikingly, treatment with JZL184 increased immobility time for both WT and HD mice, such that the genotype difference was preserved (treatment: *F*(1, 33) = 9.8, *p* = 0.004; interaction: F(1, 33) = 1.52, p = 0.23) (Fig. 3A); the result for WT mice is similar to findings of a previous study (Pavón et al., 2021). We also measured the activity of mice in an open field over a 10 min period. We found no genotype differences in either distance traveled (ANOVA, genotype: F(1, 21) = 0.04, p = 0.84) (Fig. 3B) or time spent in the center of the arena (ANOVA, genotype: F(1, 21) = 0.40, p = 0.54) (Fig. 3C), and JZL184 treatment also had no effect on locomotor activity in this task (ANOVA, distance traveled treatment: F(1, 21) = 1.09, p = 0.31; center time treatment: F(1,21) = 0.83, p = 0.37). Although YAC128 mice were heavier on average than WT mice (RM ANOVA, genotype: F(1, 34) = 17.22, p = 0.0002), there was no significant difference in weight within genotype for treated and untreated mice (RM ANOVA, WT treatment: F(1, 16) = 0.90, p = 0.36); YAC128 treatment: F(1, 18) = 0.80, p = 0.38) (Fig. 3D).

**Figure 3.**
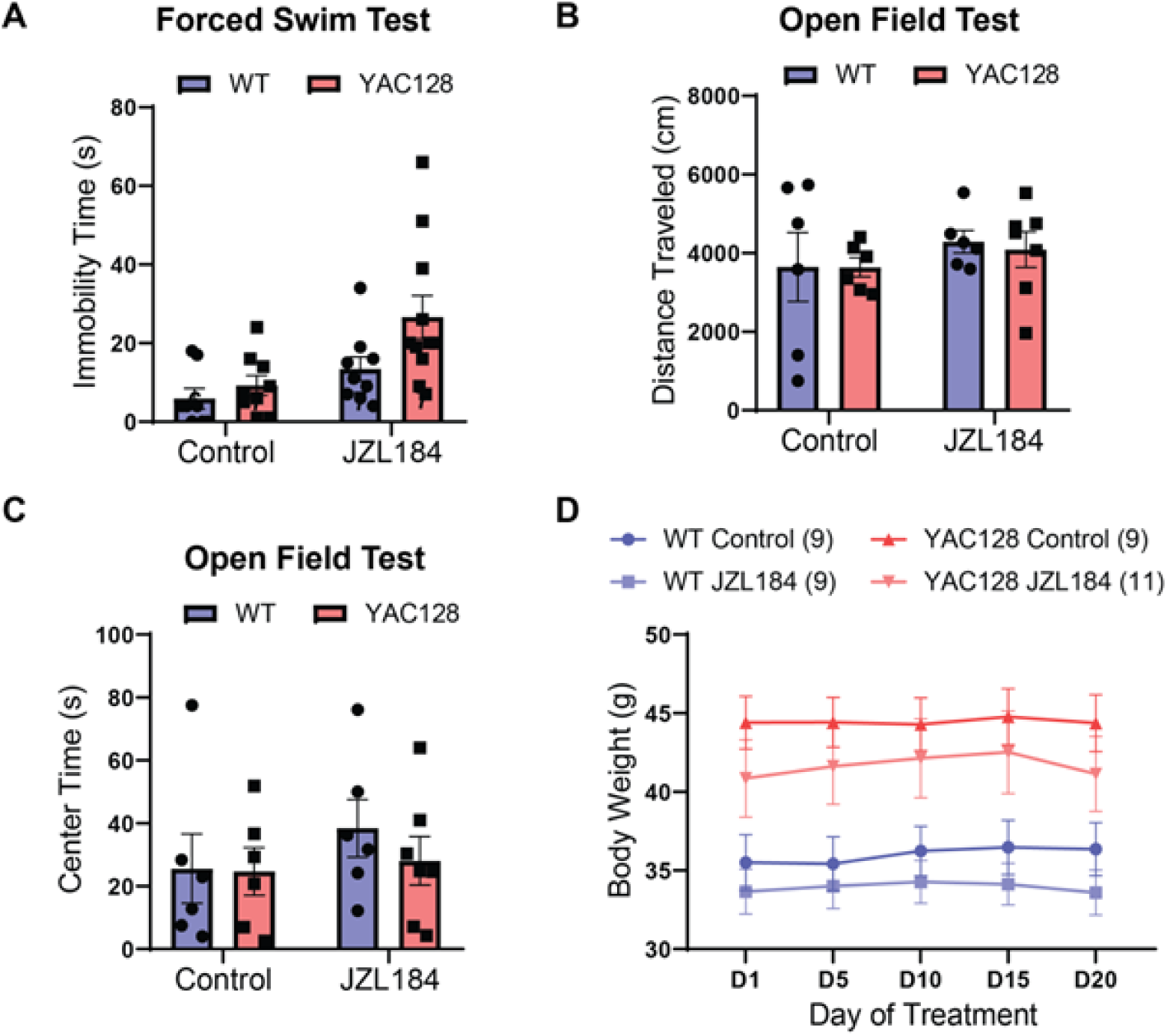
Effect of chronic JZL184 treatment on locomotor and affective behaviors. A) JZL184 treatment increased immobility time on the forced swim test across genotypes (Sidak test, p = 0.004). B) Locomotor activity, as measured by distance traveled in the open field test, was not significantly different between genotypes or treatment groups. C) Center time in the open field arena was similar across genotypes and treatment conditions. D) Bodyweight was significantly higher across treatment in YAC128 as compared to WT mice, but no effect of treatment was observed.

### JZL184 treatment enhances LTP in WT and restores synaptic plasticity at YAC128 corticostriatal synapses to WT levels

Previously, we reported that HFS-LTD was robust in brain slices from WT mice, but greatly attenuated or absent in slices from YAC128 mice (Sepers et al., 2018); notably, those experiments were from mice who had not undergone any behavioral testing (naive). Here, we used field recordings to measure the striatal neuronal population response to 100Hz stimulation of glutamatergic terminals in the striatum for 1s, repeated four times at 10s intervals (HFS). Surprisingly, the results from all four groups shown in a cumulative probability plot (Fig. 4D) demonstrate a right shift towards LTP displayed by both WT and YAC128 treated with JZL184 (RM ANOVA, F(1.5, 21) = 12.68, p = 0.0006). Moreover, instead of exhibiting LTD, as had been observed previously in naive mice, WT mice treated with vehicle that underwent behavioral testing showed slight long-term potentiation (LTP) that was significantly augmented by treatment with JZL184 (Fig. 4A,C, Sidak p = 0.004). As previously found for naive YAC128 mice (Sepers et al., 2018), vehicle-treated YAC128 mice showed no plasticity response to the HFS protocol. In contrast, JZL184-treated YAC128 mice exhibited LTP (YAC128 vs YAC128 JZL, Sidak p = 0.0110) that was indistinguishable from that found in JZL184-treated WT mice (Fig. 4B,C Sidak p = 0.1940).

**Figure 4.**
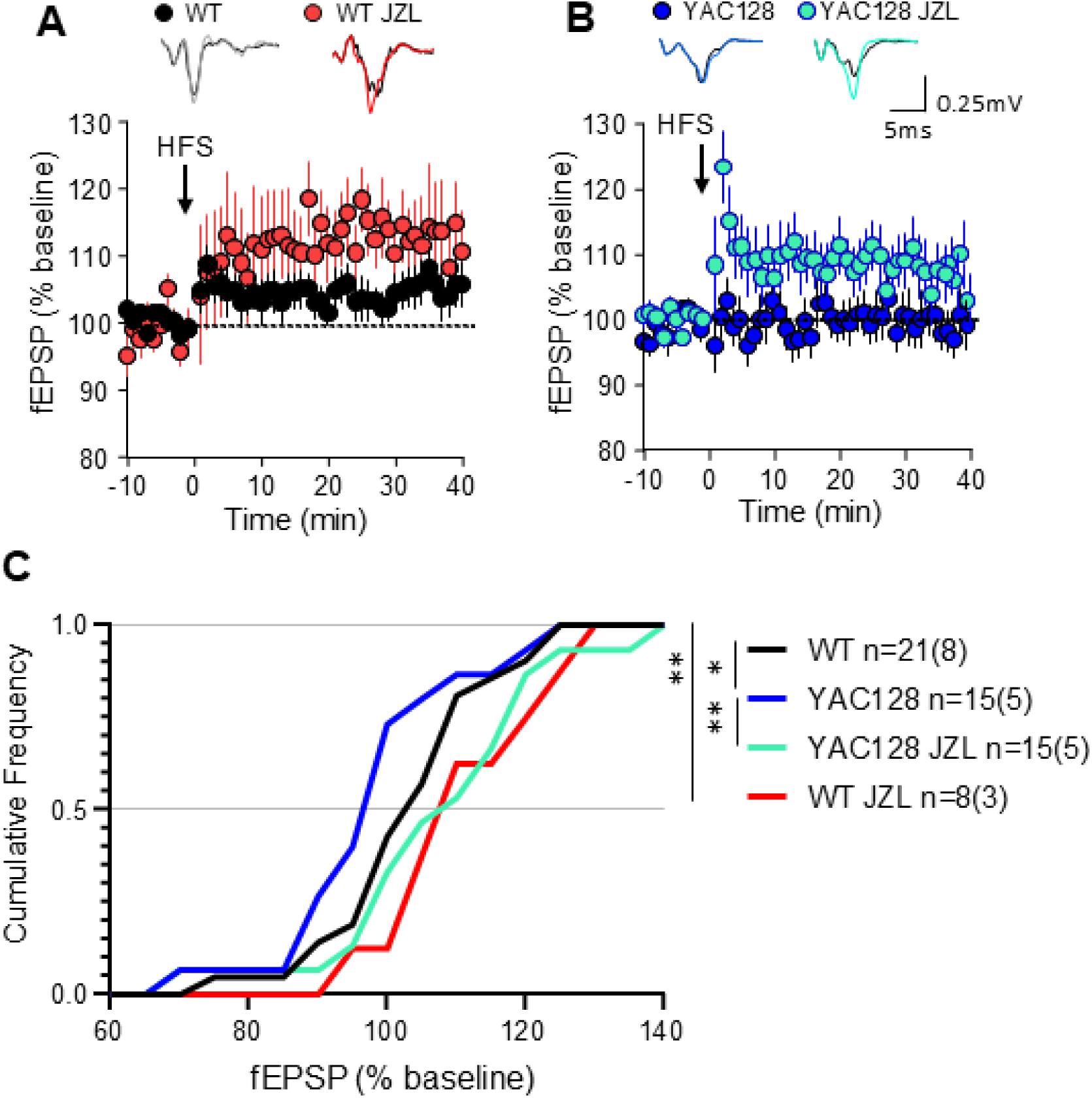
Cortico-striatal plasticity in acute brain slices following JZL184 treatment in vivo. A) Time course summary of change in field excitatory postsynaptic potential (fEPSP) amplitude normalized to baseline in response to high frequency stimulation (HFS) in WT vehicle-treated (WT) and WT treated with JZL184 (WT JZL) B) Time course summary of YAC128 fEPSP response to HFS in vehicle-treated (YAC128) and JZL184-treated (YAC128 JZL). Note: Long term depression (LTD) was not observed for any group. n = slices (mice) as shown in C for all groups. A) and B) Representative traces at top show baseline fEPSP in black C) Cumulative frequency of response size 30-35 min post HFS showed a significant shift towards increased amplitude of responses following JZL treatment in vivo in both WT and YAC128 mice. n = slices (mice) shown in C for all groups, *p<0.05, **p<0.01 by repeated measures ANOVA with Sidak multiple comparisons.

It was surprising to us to find that WT vehicle-treated mice lacked HFS-LTD, which we had previously documented in WT FVB/N mice at both 2- and 6-months of age, and instead observed a small LTP in response to HFS in striatum. We hypothesized that the shift from LTD to LTP could be a result of either: i) exposure to peanut butter (PB) in their diet +/- handling by the experimenter, or ii) a response to the behavioral testing. To further investigate this unexpected form of LTP in WT, we repeated the slice experiments in a group of naive WT mice (no handling, PB, or behavioral testing, for comparison to previous data in (Sepers et al., 2018) and in a group of mice that were handled and given PB daily for 14-19 days without any behavioral testing (PB only). We compared results to those of WT mice treated with JZL184 or vehicle (PB) that underwent behavioral testing (the same two groups of WT mice shown in Figure 4). As shown in Figure 5, the striatum of naive WT mice exhibited HFS-induced LTD and a significant increase in paired pulse ratio (PPR, paired t-test, p=0.0211) suggesting a reduction in the presynaptic probability of glutamate release, consistent with previous findings(Sepers et al., 2018). Interestingly, experiments in naive mice could be categorized as either LTD or failed plasticity without any experiments showing LTP, while PB-only mice (subjected to handling but no behavioral testing) showed a mix of responses to HFS. There was a further increase in the proportion of experiments that showed LTP in the behaviourally-tested vehicle- and JZL184-treated WT mice. Moreover, the PPR in vehicle-treated WT mice that underwent behavioral testing was not changed by HFS (paired t-test, p=0.5636). Comparison of the distribution of HFS responses showed that behaviourally-tested WT (same as “WT vehicle” in Fig. 4) were significantly shifted towards LTP compared to naive mice and PB alone (RM ANOVA, F(1.3,18.4) = 18.41, p=0.0002), suggesting that this metaplasticity is experience dependent.

**Figure 5.**
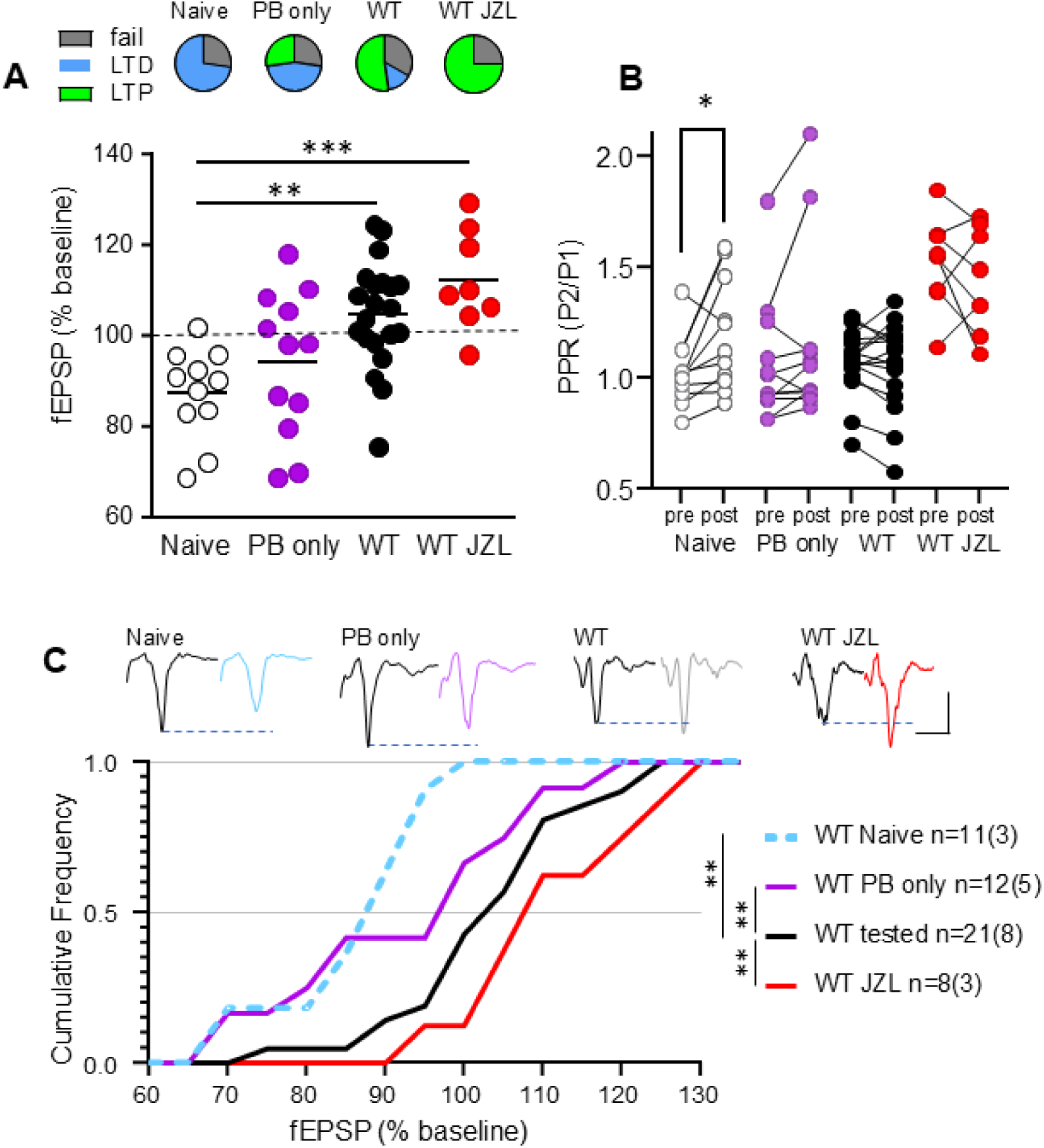
In WT mice, behavioral testing and JZL treatment change striatal synaptic plasticity. “Naive” WT were not handled, exposed to peanut butter or to behavioral testing before the striatum was collected for experiments. WT “PB only” were handled daily and given peanut butter (vehicle) for 3 weeks without any behavioral testing. The WT group was the original vehicle-treated control and were given PB and exposed to behavioral testing at the same time as the WT JZL group. Note: WT and WT JZL group data are also shown in figure 4 but copied here for comparison with PB only and naive groups. A) Top panel: Proportion of experiments with fEPSP amplitude after HFS that showed no change (fail), a significant decrease (LTD) or an increase (LTP). Bottom panel: Summary of change in fEPSP in response to HFS in each group where the Naive WT only showed LTD or fail, in contrast to the behaviorally tested and treated groups that all showed some experiments with LTP. ANOVA with Sidak post hoc.B) Paired-pulse ratio of fEPSP (PPR) shows a significant increase after HFS (post) compared to baseline (pre) only in Naive WT striatum (paired t-test). C) Cumulative frequency of response size 30-35 min post HFS showed a significant difference between groups. RM ANOVA with Sidak multiple comparisons. \**p*<0.05, \*\**p*<0.01, \*\*\**p*<0.001.

## Discussion

Although the endocannabinoid system is ubiquitous throughout the brain and involved in many forms of memory (Martinez Ramirez et al., 2023), endocannabinoid-dependent corticostriatal synaptic plasticity has been specifically implicated in skilled motor learning in rodents (Lovinger, 2010). We previously reported a deficit in one form of this plasticity – HFS-LTD – in acute brain slices from YAC128 mice (Sepers et al., 2018). That plasticity deficit could be normalized to WT levels by perfusing slices with JZL184, an inhibitor of monoacylglycerol lipase – the enzyme responsible for degrading 2-AG. Here, we have shown that chronic administration of JZL184, which we have demonstrated increases basal striatal levels of 2-AG by >5-fold, restores corticostriatal synaptic plasticity and also improves rotarod motor learning in 5-6m old YAC128 mice, matching that of WT littermates.

### eCBs and corticostriatal plasticity in motor learning

Dynamic changes in connections between dorsal striatum and prefrontal and sensorimotor cortex are involved in motor learning in rodents and humans (Balleine and O’Doherty, 2010; Costa et al., 2004; Dayan and Cohen, 2011). Synapses between cortical glutamatergic projections that drive activity of striatal GABAergic SPNs show plasticity in response to trains of stimuli (as recorded in *ex vivo* brain slice from untrained animals) that is postulated to underlie new motor learning (Balleine and O’Doherty, 2010; Graybiel and Grafton, 2015; Klaus et al., 2019; Perrin and Venance, 2019). While one *in vivo* study in WT mice showed that temporal changes in cortical terminal Ca^2+^ dynamics in dorsal striatum are linked to stages of learning the accelerating rotarod task (Kupferschmidt et al., 2017), the underlying mechanisms of these changes remain unknown. eCBs are lipid-based neurotransmitters that suppress Ca^2+^ influx and thereby reduce neurotransmitter release at axon terminals and mediate synaptic plasticity recorded from *ex vivo* brain slices (Kreitzer, 2005; Lovinger, 2010). eCBs have been implicated in gating the transition from goal-directed to skill consolidation (habit formation) in motor actions, which is critical for learning and flexibility in executing skilled movement (Gremel et al., 2016). Moreover, 2-AG has been shown to be involved in a reward-based motor sequencing task (Tanigami et al., 2019), and a deficit in eCB-mediated corticostriatal synaptic plasticity alters motor learning (Benthall et al., 2022). Acute pharmacological activation or inhibition of CB1R have both been linked to impaired motor learning, whereas chronic CB1R agonism can reinforce habit learning mediated by corticostriatal connections (Goodman and Packard, 2015; Mato et al., 2004). Given these previous studies, our results — showing that chronic JZL184 treatment to elevate basal and on-demand levels of 2-AG restores motor learning on the rotarod in YAC128 mice – are consistent with our previous finding that JZL184 also normalizes CB1R-mediated HFS-LTD in cortical-striatal brain slices from this HD mouse model (Sepers et al., 2018).

Our previous results are consistent with many studies showing that corticostriatal synapses *ex vivo* express LTD in response to a variety of stimulus protocols (Lovinger, 2010; Wang et al., 2006). In contrast, LTP is only observed with complex spike timing protocols or theta burst stimulation and differs between dSPN and iSPN (Hawes et al., 2013; Perrin and Venance, 2019; Shen et al., 2008). CB1R activation is well known to reduce glutamate release to induce LTD, but eCBs can mediate bidirectional plasticity *in vivo* (Xu et al., 2018). Although rarely studied, *ex vivo* striatal plasticity can be altered by *in vivo* experiences such as motor learning or environmental enrichment (Hawes et al., 2015; Morera-Herreras et al., 2019; Yin et al., 2009).

Here we show that synaptic plasticity evoked in the striatum is changed in vehicle-treated WT mice after behavioral testing compared to naive WT mice. The greater shift in plasticity shown after JZL184 treatment suggests an *in vivo* change that biases responses to HFS towards potentiation instead of depression of excitatory synapses in the striatum. Future studies are needed to determine the mechanism of this effect.

### CB1R expression and corticostriatal connectivity changes in Huntington’s disease

Several studies have reported reduced CB1R in the striatum at early motor manifest stages of HD. Positron Emission Tomography imaging with CB1R ligands shows significantly reduced signal in striatum in patients with Stage 1 HD (Van Laere et al., 2010), and HD mouse models exhibit a reduction in CB1R transcription and protein levels in pre-motor manifest stages (Blázquez et al., 2011; Dowie et al., 2009; Glass et al., 2000; Mievis et al., 2011). Notably, the early reduction in CB1R in the striatum of mouse models occurs in SPNs (Denovan-Wright and Robertson, 2000), while CB1R is preserved on cortical terminals within the striatum until late stages (Chiodi et al., 2012). Moreover, selective knock-out of CB1R on these glutamatergic terminals in striatum impairs rotarod performance in WT mice and accelerates the HD phenotype in R6/2 mice, whereas selective knock-out in GABAergic striatal neurons has no effect on R6/2 phenotype or on WT rotarod performance (Chiarlone et al., 2014). Consistent with those data, genetic rescue of CB1R expression in striatal SPNs of R6/2 mice did not improve rotarod performance, although there was an indirect effect to preserve cortical glutamatergic connections with striatal SPNs (Naydenov et al., 2014). Our recent results in *ex vivo* brain slice of YAC128 mice also show intact CB1R expression and function on cortical terminals in striatum at a time when corticostriatal HFS-LTD is impaired due to a deficiency of on-demand synthesis of anandamide in SPNs(Sepers et al., 2018). In the present study, we show no genotype difference in tonic, baseline 2-AG or anandamide levels, suggesting that the deficit in eCB-mediated HFS-LTD in the YAC128 mice is a result of impaired signaling to induce dynamic, phasic increases in eCB levels upon high-frequency stimulation. The deficit in HFS-LTD progresses to complete loss by 6 months of age in YAC128 mice (Sepers et al., 2018). Rotarod learning is impaired by 4 months of age in YAC128 mice (Milnerwood et al., 2010), while loss of glutamatergic synapses on SPNs occurs only later, close to 12 months of age (Wu et al., 2016). Taken together, results of those studies suggest early impairment in rotarod learning is not because of a cortical-striatal disconnection, or reduced CB1R on cortical terminals; instead, those studies, together with the data presented here showing normalization of rotarod learning in 6-month YAC128 mice treated with JZL184 to boost eCB levels, argue for an early deficit in cortical activity-induced eCB synthesis in SPNs as a major contributing factor to reduced motor learning in HD mice.

### Effect of JZL184 on immobility in forced swim test

Although immobility time in the FST has been used in the past as a measure of depressive mood in rodents, more recent studies suggest it more directly reflects response to stress and is correlated with dopamine levels in prefrontal cortex (Pavón et al., 2021). Notably, immobility time was increased in both WT and YAC128 mice treated with JZL184; therefore, it is possible that one down-side of chronic treatment to elevate eCB levels is to promote passive avoidance responses to stress rather than active effort to reduce exposure to the stressful stimulus/environment. Alternatively, the reduced swim time may reflect an overall effect of elevated 2-AG levels to reduce intense motor activity, which would not be required for activity in the open field testing. Chronic JZL184 has been tested in vivo at doses ranging from 4 - 40mg/kg/day in mice (Ghosh et al., 2015; Kinsey et al., 2013; Schlosburg et al., 2010) and higher doses lead to tolerance and desensitization of CB1R. In contrast, a lower dose of 4mg/kg/day maintained normal CB1R expression and synaptic function. Although FST was not assessed in those studies, the lower dose didn’t impact open field behaviour (Ghosh et al., 2015) consistent with our results.

### Modulating CB1R in patients with Huntington’s disease

To date there have been only a few studies modulating the endocannabinoid system in patients with HD, despite the literature supporting the important role of eCBs in motor learning and control. These have been small trials of open-label or placebo-controlled treatment with direct CB1R agonists, and results show little impact on motor control, cognitive deficits or psychiatric disturbances (Curtis et al., 2009; López-Sendón Moreno et al., 2016). Treatment with a compound that instead boosts levels of eCBs on demand, as required to mediate forms of activity-dependent synaptic plasticity underlying motor learning, may be more effective with potentially fewer side effects.

## Acknowledgments and Funding

We are grateful to Lily Zhang for help with animal care and genotyping, as well as assistance with administration of peanut butter treatment. This work was supported by grants from the Huntington Society of Canada and the Canadian Institutes of Health Research (CIHR) PJT-178043 to LAR. CLW was supported by a Canada Graduate Scholarship - Doctoral award. LAR holds the Louise A. Brown Chair in Neuroscience. MNH was supported by operating funds from CIHR. HAV was supported by a scholarship from Branch Out Neurological Foundation.

## Author Contributions

CLW, MDS, LAR and MNH designed the experiments. CLW, MDS, DR, and HAV performed the research. CLW and MDS analyzed the data. CLW, MDS and LAR wrote the manuscript with input from DR and MNH.

